# Characterisation of antigenic MHC Class I-restricted T cell epitopes in the glycoprotein of Ebolavirus

**DOI:** 10.1101/494021

**Authors:** Jonathan Powlson, Daniel Wright, Antra Zeltina, Mark Giza, Morten Nielsen, Tommy Rampling, Navin Venkatrakaman, Thomas A. Bowden, Adrian V.S. Hill, Katie J. Ewer

**Affiliations:** The Jenner Institute, Old Road Campus Research Building, University of Oxford, Oxford, UK.; Division of Structural Biology, Wellcome Centre for Human Genetics, University of Oxford, Oxford, UK.; Department of Bio and Health Informatics, The Technical University of Denmark, Lyngby, Denmark.

## Abstract

Ebolavirus is a pathogen capable of causing highly lethal haemorrhagic fever in humans. The envelope-displayed viral glycoprotein is the primary target of humoral immunity induced by both natural exposure and vaccination. The epitopes targeted by B cells have been thoroughly characterised by functional and structural analyses of the glycoprotein, GP, yet there is a paucity of information regarding the cellular immune response to Ebolavirus. To date, no T cell epitopes in the glycoprotein have been characterised in detail in humans.

A recent Phase I clinical trial of a heterologous prime-boost vaccination regime with viral vectors encoding filovirus antigens elicited strong humoral and T cell responses in vaccinees. Using samples from this trial, the most frequently recognised peptide pools were studied in more detail to identify the minimal epitopes recognised by antigen-specific T cells and associated HLA-restrictions.

Using IFNγ ELISPOT and flow cytometry, we characterised nine highly immunogenic T cell epitopes located on both the GP_1_ and GP2 subunits of the Ebolavirus GP. Epitope mapping revealed the location of these epitopes as presented on the mature virion. HLA-typing on all participants, combined with *in silico* epitope analysis, determined the likely MHC class I restriction elements. Thirteen HLA-A and -B alleles were predicted to present the identified epitopes, suggesting promiscuous recognition and induction of a broad immune response.

The glycoprotein of Ebolavirus is highly immunogenic, inducing both CD4^+^ and CD8^+^ T cell responses and we have shown here for the first time that these responses are associated with multiple HLA types. Delivery of this antigen using a heterologous prime-boost approach with ChAd3 and MVA is likely to be highly immunogenic in genetically diverse human populations, due to the induction of responses against multiple immunodominant epitopes.

**Author Summary:** The West African Ebolavirus epidemic in 2013-2016 claimed more than 11,000 lives, eclipsing all previously recorded outbreaks combined. Understanding the mechanisms of host-pathogen interactions informs rational vaccine design. B cell epitopes have been described extensively through the characterisation of monoclonal antibodies from survivors of Ebolavirus disease. Despite a clear role for cellular immunity in survival of Ebolavirus disease, data relating to minimal T cell epitopes in the glycoprotein has so far only been obtained in murine. This study describes the first identification of human T cell epitopes in the surface glycoprotein of the Ebolavirus. These epitopes have been identified using samples from a clinical trial of viral vectored vaccines against the Zaire Ebolavirus undertaken during the 2013-2016 West African outbreak. We also describe the recognition of these epitopes through multiple HLA alleles. As efforts to progress a vaccine against Ebola towards licensure proceed, detailed characterisation of immunity to the virus contribute to our understanding of the potential application of vaccines in diverse populations.

## Introduction

The West African Ebolavirus epidemic that began in Guinea in December 2013 claimed more than 11,000 lives, eclipsing all previously recorded outbreaks combined [1, 2]. The aetiological agent of this outbreak, *Zaire ebolavirus* (EBOV), is one of five species within the *Ebolavirus* genus of the *Filoviridae* family [3]. Three other members of this genus; *Sudan ebolavirus* (SUDV), *Tai Forest ebolavirus* (TAFV) and *Bundibugyo ebolavirus* (BDBV), are also pathogenic in humans [4]. The fifth member, *Reston ebolavirus* (RESTV), is not associated with disease in humans but has been isolated from non-human primates (NHP) and swine [5, 6]. While the case fatality rate of the West African epidemic (2013-2016) was estimated at 40%, it has been as high as 90% in previous outbreaks [1]. Despite the threat that EBOV and other filoviruses pose, relatively little is known about the cellular immune response to filoviruses in humans.

Rodent and NHP models, in addition to data from survivors of Ebolavirus disease, have demonstrated the importance of both humoral and cellular responses in clinical outcomes of EBOV disease [7–11]. Both T cell and antibody responses to EBOV are directed against the viral glycoprotein (GP) [8, 12], which is essential for attachment, fusion and entry of the virion into the target cell [13]. The GP protein is synthesised as a 676-amino acid polypeptide which undergoes post-translational cleavage by the host cell proprotein convertase furin. This cleavage yields two disulphide-linked subunits, GP1 and GP2, which further trimerise to form a ‘chalice-like’ structure [14]. The membrane-distal GP_1_ displays both N- and O-linked glycosylation and is responsible for host cell attachment. The smaller GP_2_ fragment anchors the complex to the envelope via a transmembrane domain and contains a hydrophobic internal fusion loop which drives fusion of the host and virion membranes, facilitating the release of viral nuclear material into the cytosol [13].

The efficacy of monoclonal antibody therapies targeting the glycoprotein has been demonstrated *in vivo* in guinea pig and macaque models with protection seen up to 5 days post-challenge when administered as either a monotherapy or monoclonal or polyclonal cocktail [15–18]. Furthermore, the experimental human vaccine rVSV-ZEBOV, which showed 100% efficacy in a ring-vaccination study in Guinea, appears to protect via a predominantly antibody-mediated mechanism [19–21], particularly given the paucity of T cell responses to the vaccine [12, 22–24]. Whilst solid progress has been made understanding humoral immune responses to EBOV in humans, less is known about the mechanisms underlying cellular immunity. It has been clearly demonstrated that CD8^+^ cytotoxic T cells (CTL) are required for survival in macaques and that depletion of this subset in vaccinated animals results in complete abrogation of protection in lethal challenge models [11, 25–30]. T cell epitopes in EBOV nucleoprotein (NP) have been identified in immunized mice and several computational studies have predicted further epitopes in both NP and GP. Although EBOV T cell epitopes in the nucleoprotein have been characterised in human studies of Ebola survivors, to date no minimal epitopes have been characterised in the glycoprotein [31]. This is despite the potent immunogenicity of the glycoprotein both in the context of natural infection [8, 9] and in vaccines expressing GP as the target antigen [22, 32–34].

Identification of immunogenic epitopes in pathogen genomes has been used as a tool to generate novel vaccine candidates and identify potentially protective targets for CTL [35]. A number of *in silico* bioinformatics tools are available for prediction of epitopes and associated HLA-restriction elements [36, 37]. This information can be used to design so-called “mosaic” antigens; an approach that has been widely used for the development of novel HIV vaccines [38–40], particularly where broad coverage of conserved proteins is required. Characterisation of protective HLA-restricted CD8^+^ T cell epitopes has been undertaken for a range of diseases including influenza A[41, 42]. This information can be used to study mechanisms of immunity to inform rational vaccine design [43].

Heterologous prime-boost vaccination with viral vectors elicits strong antibody and CTL responses making them ideal candidates for immunisation against EBOV. We have previously reported a clinical trial in which volunteers were primed with a monovalent chimpanzee adenovirus encoding the EBOV GP gene [33]. One to ten weeks later, individuals were boosted with the multivalent MVA-BN-Filo, which encodes the EBOV, SUDV and *Marburgvirus* GP and TAFV NP genes. Peripheral blood mononuclear cells (PBMC) from these individuals were used to identify thirteen highly reactive nonamers. Mapping analysis revealed nine spatially distinct epitopes on the surface of EBOV GP, providing a detailed characterization of the peptides responsible for initiating CTL-mediated responses to EBOV infection.

## Results

### ELISpot responses to pooled peptides

As part of the immunomonitoring performed during the previously reported clinical trial [33], freshly isolated PBMC were assayed against 10 pools of overlapping peptides spanning the length of the vaccine antigen insert, which includes a signal peptide and the GP_1_ and GP2 subunits. Details of the peptide pools are given in Supplementary Table 1. Seven peptide pools were highly immunogenic (GP 1.1, 1.2, 1.3, 1.4, 1.6, 2.1 and 2.2) and four of the pools gave geometric mean (GM) responses greater than 150 spot-forming cells per million PBMC (SFC) across the 28 volunteers tested. These were pools 1.1 – 1.3 and 2.2 (Figure 1A). The strongest responses were seen to pools 1.1 and 2.2, corresponding to the N-terminus of GP_1_ and C-terminus of GP2, with GM responses of 237 and 164 SFC respectively. The peptides constituting the signal peptide (SP), as well as 1.5 and 1.7 pools gave positive responses in less than 40% of participants and were not assessed further.

**Figure 1.**
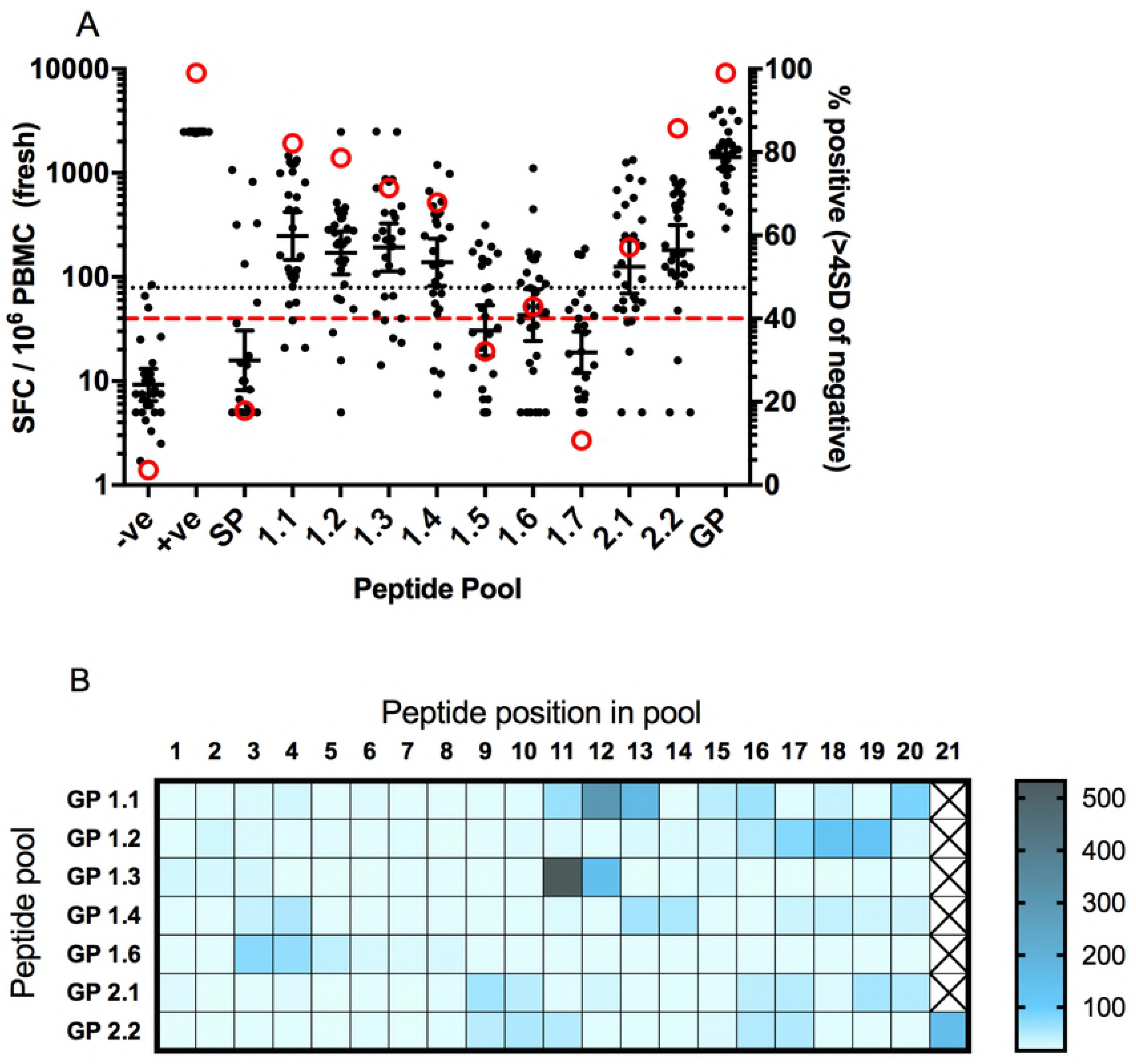
Ex-vivo IFNγ ELISPOT responses to peptide pools and 15mer peptides in individual participants. A. Responses to peptide pools using freshly isolated PBMC at seven days after boosting. Black dots represent individual participants (n=28), bars show geometric mean with 95% CI. Dotted line shows lower limit of detection for the ELISPOT assay (LLD). Red circles show the percentage of participants with a positive response to each pool. Red dashed line indicated threshold used to select pools for further analysis (78.2 SFC, equivalent to 4SD from the median of the negative control responses).-ve, negative control (medium only); +ve, positive control; GP, 2 pools containing all 187 peptides. B. Heatmap showing responses to individual peptides for each pool. Geometric mean responses are displayed (n=6-8 participants per pool, depending on sample availability).

### ELISpot responses to individual peptides

The seven most immunogenic pools were retested by IFNγ ELISpot using frozen PBMC selected from clinical trial participants that responded strongly in the preliminary assay, subject to sample availability. Each pool was deconvoluted to the constituent peptides (20-21 peptides per pool) using PBMC from between 6 and 8 vaccinees. In the seven pools tested, at least one immunodominant peptide was identified in each, and in some pools, more than one immunogenic region was observed suggesting multiple epitopes (Figure 1B and Figure 2). Following deconvolution of the responses to the peptide pools, ten of the most immunodominant peptides and regions were selected based on the strength of the ELISPOT response and sample availability, and a library of 94 nonamers was produced. A summary of the nonamers is shown in Table 1 and sequences for the individual nonamers are shown in Supplementary Table 2.

**Table 1.**
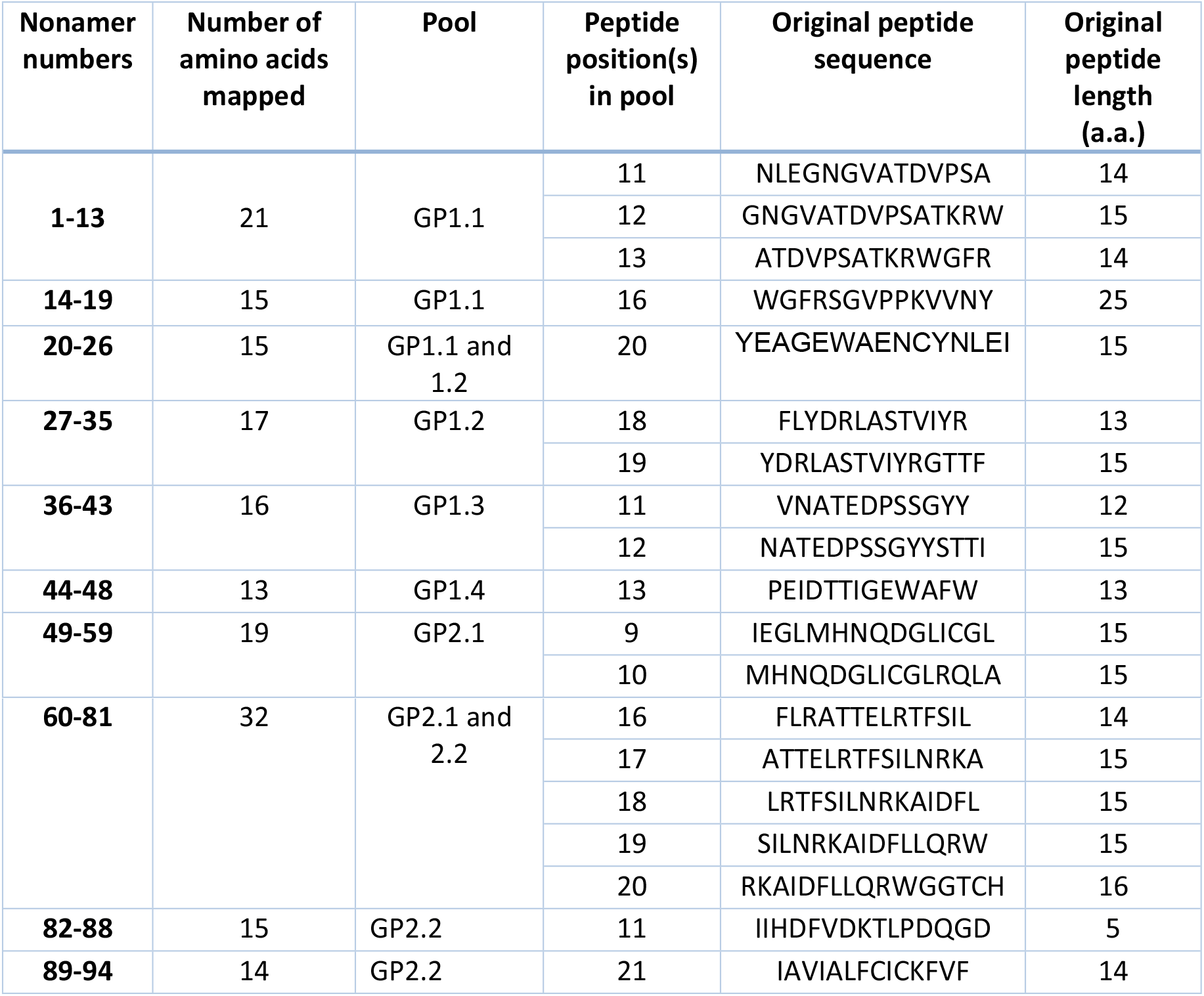
Nonamers synthesised spanning dominant 15mer peptide or regions of peptides.

**Figure 2.**
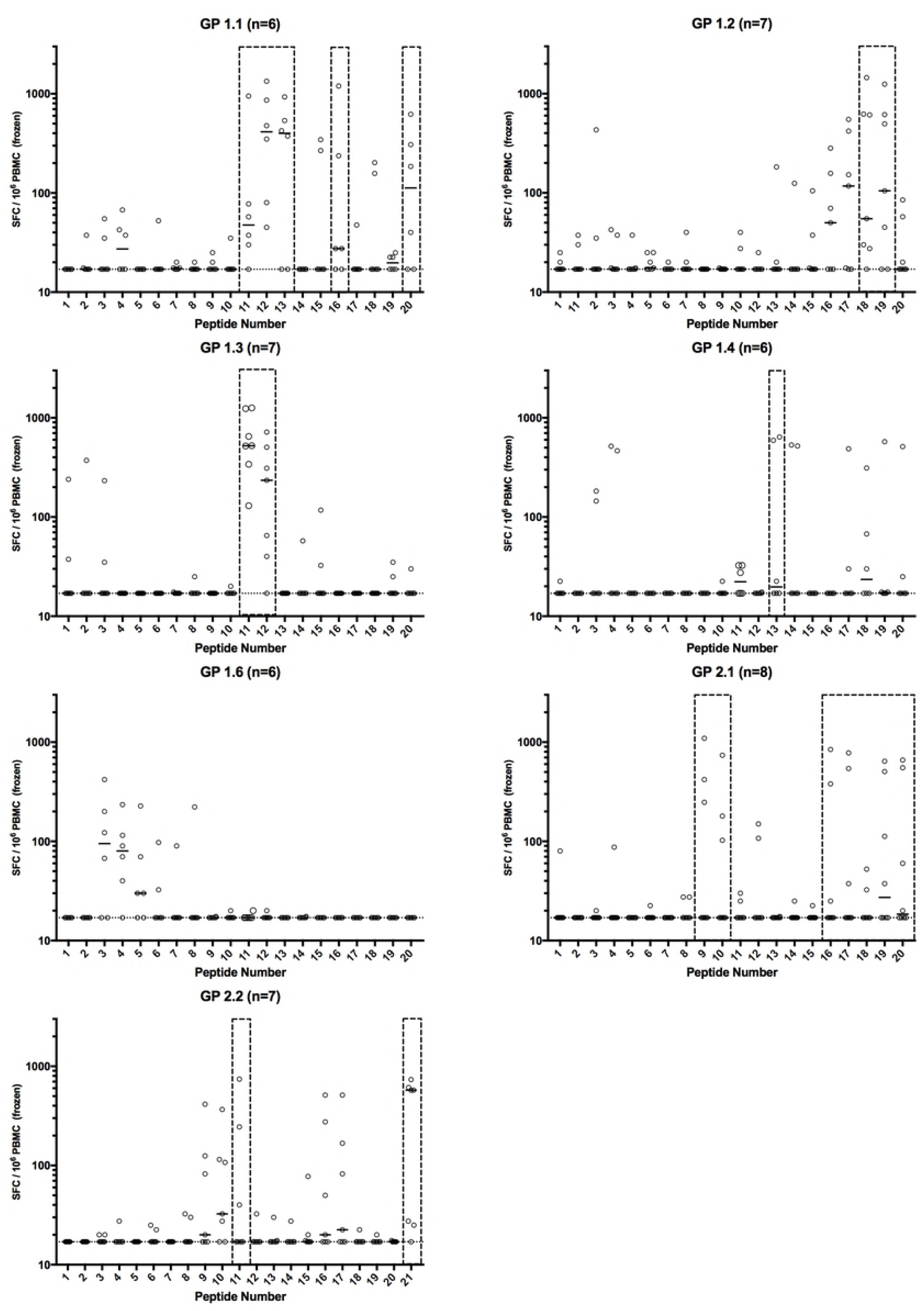
Deconvolution of individual peptide responses within peptide pools. Boxes show peptides selected for determination of minimal epitopes. Solid lines represent medians and dotted line shows lower limit of assay detection. N refers to the number of vaccinees tested for each peptide pool.

Thirteen of the nonamers elicited positive responses in more than one participant (8, 9, 15, 26, 31, 38, 39, 47,48, 55, 65, 92, 93 and 94) and responses to the nonamers are shown in Figure 3A. Where nonamers that gave a positive response were adjacent, we subsequently considered these to be part of a single epitope. These putative epitopes are displayed as nine colour coded regions. The locations of these putative epitopes were then displayed diagrammatically on the primary glycoprotein structure (Figure 3B).

**Figure 3.**
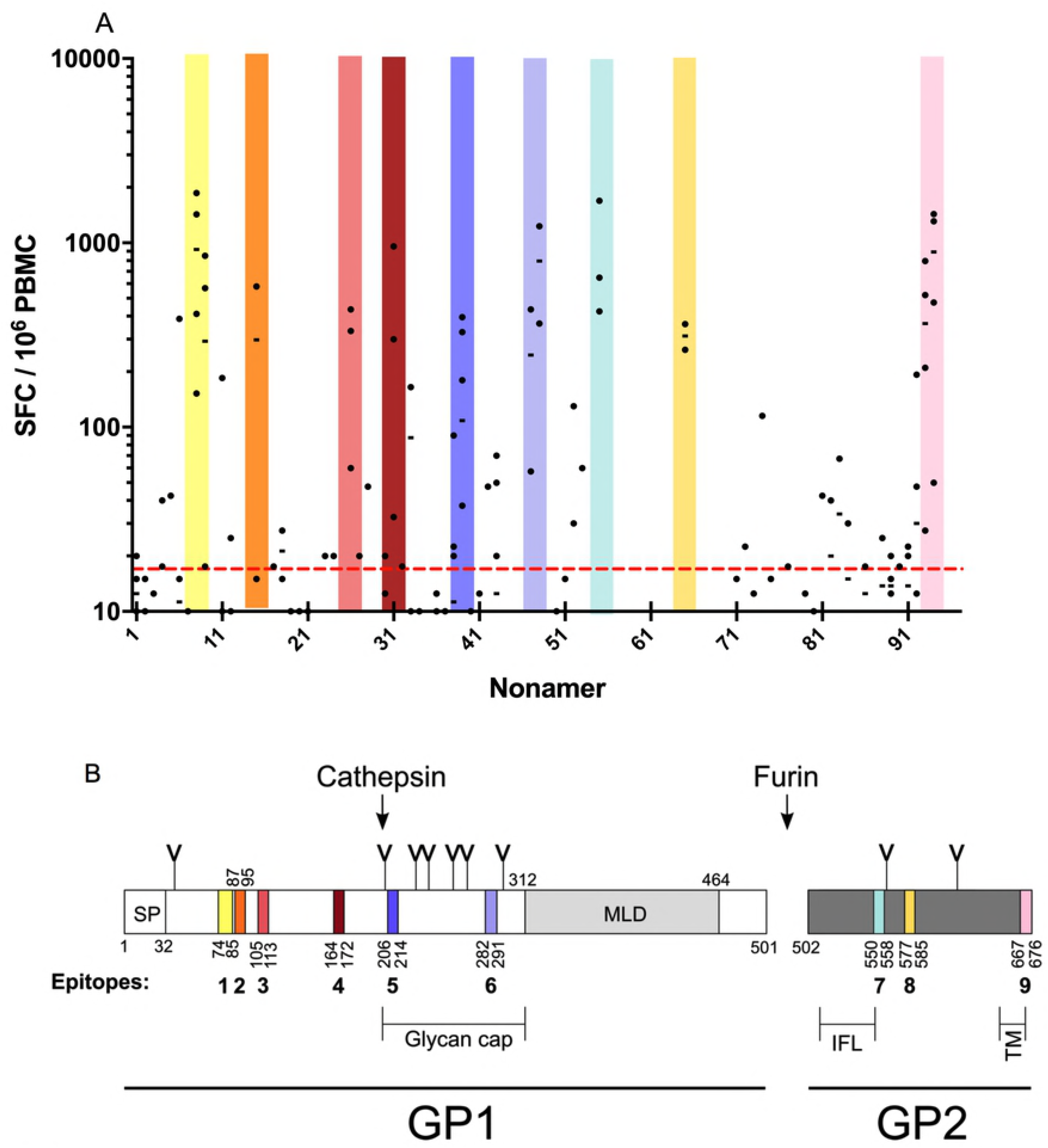
Identification of immunodominant 9mers using ex-vivo IFNγ ELISPOT. **A.** Individual 9mer peptides were tested in participants previously identified as responders to the peptide pool (n=2-6). Bars represent medians. Dashed red line indicates positivity threshold (4SD from the median of negative control). Thirteen of the nonamers elicited positive responses in more than one participant (8,9, 15, 26, 31, 38, 39, 47,48, 55, 65, 92, 93 and 94) and these are displayed as nine colour coded regions. Adjacent immunogenic nonamers that overlapped were considered to form a single epitope. B. Domain organization of the EBOV GP with the location of the T cell epitopes on the primary glycoprotein structure shown in colour. SP, signal peptide; IFL, internal fusion loop; TM, transmembrane domain; MLD, mucin-like domain; Y-shaped symbols designate N-linked glycosylation sites.

### EBOV GP Epitope Sequence and Location in Primary Structure

The 13 immunogenic nonamers formed nine discrete putative epitopes (Table 2). The sequences of the constituent nonamers were used to identify the location of the epitopes in the primary structure of EBOV GP (Figure 3B). The first three are located in an epitope-rich 40 amino acid stretch in the N-terminal half of GP_1_. The first epitope is 12 amino acids long and comprised of the nonamers 6, 8 and 9 although nonamer 7 elicited no T cell activity (Figure 3B). The second epitope is situated immediately adjacent to the first and is comprised of nonamer 15 alone. No immunogenicity was detected with nonamers 11-14, which span the junction of the two epitopes. The third epitope is located nine residues downstream. A fourth epitope, constituting nonamer 31 alone, corresponds to residues 164-172. These four epitopes are all involved in the formation of the GP_1_ head structure in the mature GP (Supplementary figure 1). Epitopes five and six are located within the glycan cap region of GP_1_. No epitopes were identified near the C-terminus of GP_1_ which corresponds to pools 1.5, 1.6 and 1.7 (pools 1.5 and 1.7 were excluded from the epitope analysis at an earlier stage due to poor immunogenicity). Further analysis of the location of the epitopes on the structure of the glycoprotein is included in the Supplementary information (Supplementary Figure 1).

**Table 2.**
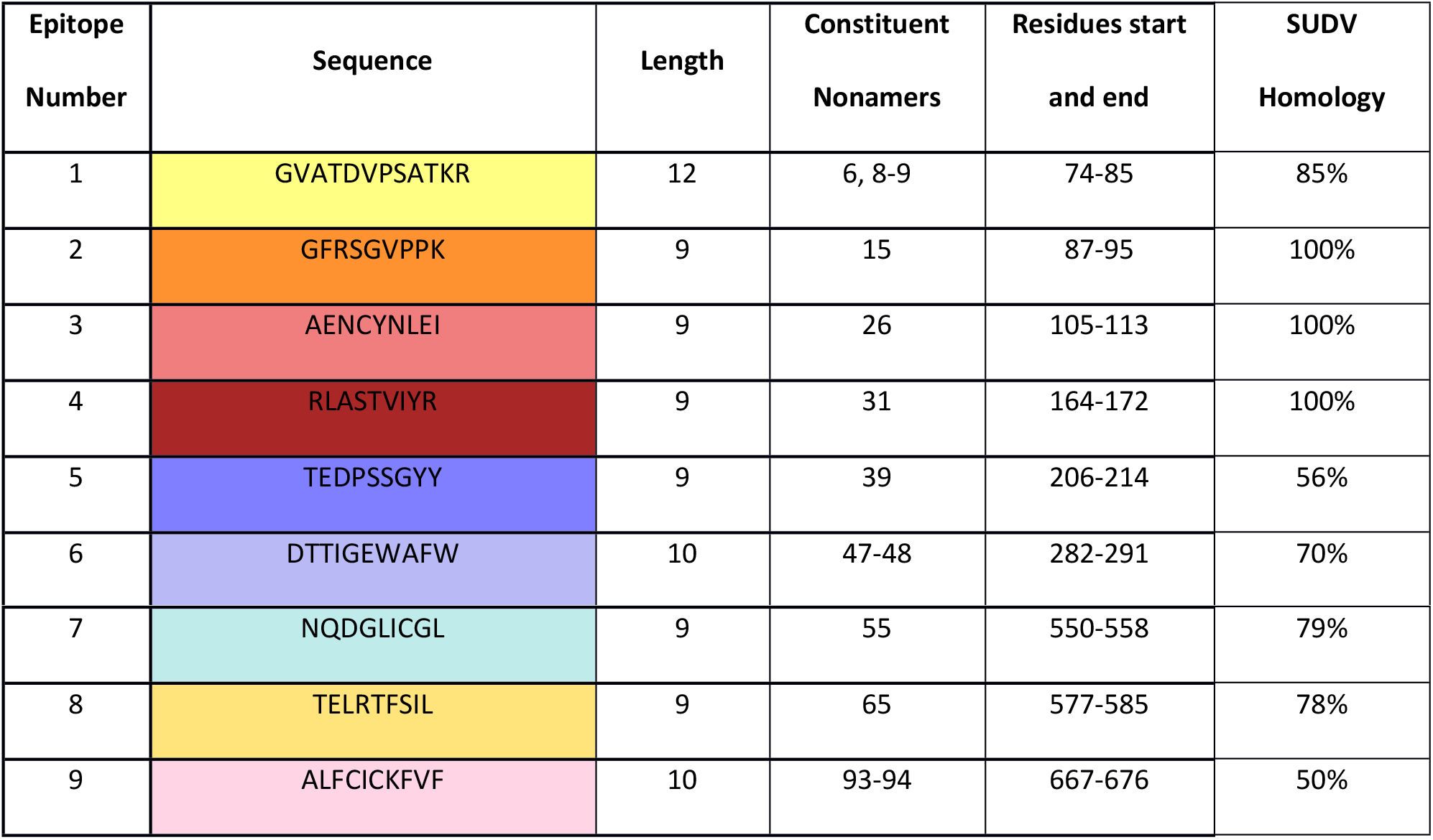
Putative epitope sequences with colour coding, residue position, constituent nonamers and SUDV homology.

We identified three epitopes in the GP_2_ protein, the first of which is 9 amino acids long formed by nonamer 55 and incorporates part of the internal fusion loop (7). The second epitope is comprised of the singular nonamer 65, and the final epitope represents the terminal ten amino acids of the GP which form part of the transmembrane domain and cytosolic tail. The other species within the genus *Ebolavirus* that causes significant outbreaks in humans is SUDV and therefore we determined sequence identity between the identified EBOV GP epitopes and the corresponding sequences in SUDV GP. Homology was at least 50% for all epitopes with epitopes 2, 3 and 4 being 100% identical (Table 2).

### CD8^+^ T cell responses to nonamers

Using flow cytometry with intracellular cytokine staining, we assayed the highest responding volunteers from the nonamer ELISpot (n=13). We determined that all eight of the nonamers tested induced IFNγ and TNFα production in CD8^+^ T cells, with the strongest responses of both cytokines induced by nonamers 8, 31 and 55. These three nonamers induced IFNγ production with a median frequency of 0.81%, 0.49% and 0.58% of CD8^+^ T cells, respectively. The median frequency of TNFα-secreting CD8^+^ T cells after stimulation with the same three nonamers was 0.74%, 0.43% and 0.62% of CD8^+^ T cells, respectively. (figure 4A).

**Figure 4.**
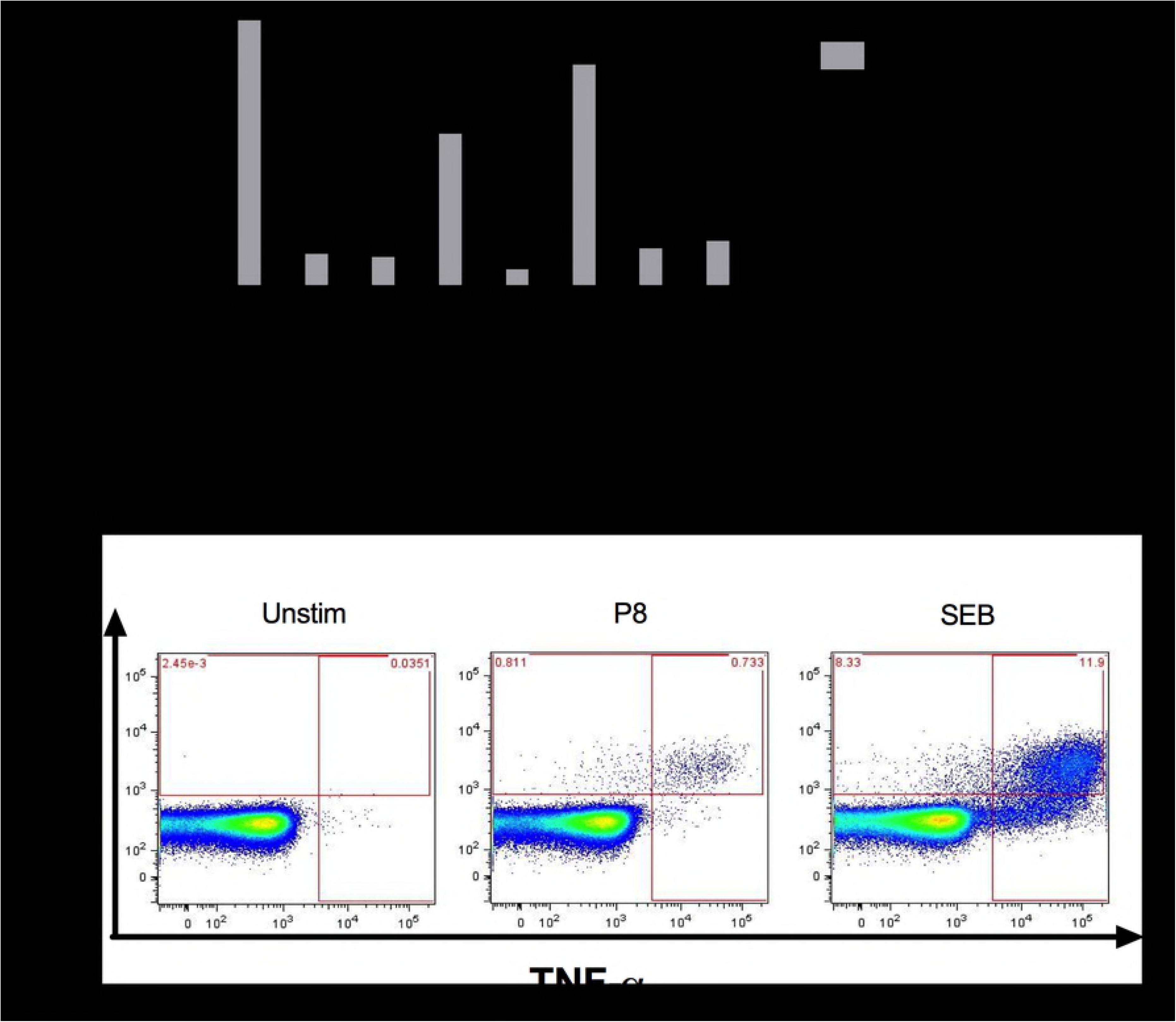
Cytokine expression from CD8^+^ T cells assessed by flow cytometry. (A) T cells producing IFNγ or TNFα as a percentage of CD8^+^ cells; bars represent single values or medians (n=1-3 per nonamer); error bars represent range. (B). Representative example of intracellular cytokine staining of CD8^+^ T cells. A full gating strategy is shown in Supplementary figure 2.

### HLA Typing and epitope restriction analysis

High-resolution HLA typing was performed on all participants (Supplementary Table 3) and MHC Class I alleles were analysed using NetMHCpan (version 3.0) to assign HLA restriction elements from each volunteer to each of the positive peptides. This predicts the binding of the peptide to each of the 3-6 HLA alleles for the volunteer, (2 each of HLA-A, B and C alleles, unless homozygous) and the allele with the strongest predicted binding value as the likely restriction element was determined. A rank score binding value of less than two percent has been shown to recover 95% of previously validated epitopes with higher than 99% specificity. For all epitopes, at least one restriction element from a responding participant was predicted to be a strong binder (Table 3) and for four of the epitopes (1, 3, 7 and 9), more than one element was predicted to bind strongly. Of the alleles predicted to bind strongly, nine were HLA-A alleles and five were HLA-B.

**Table 3.**
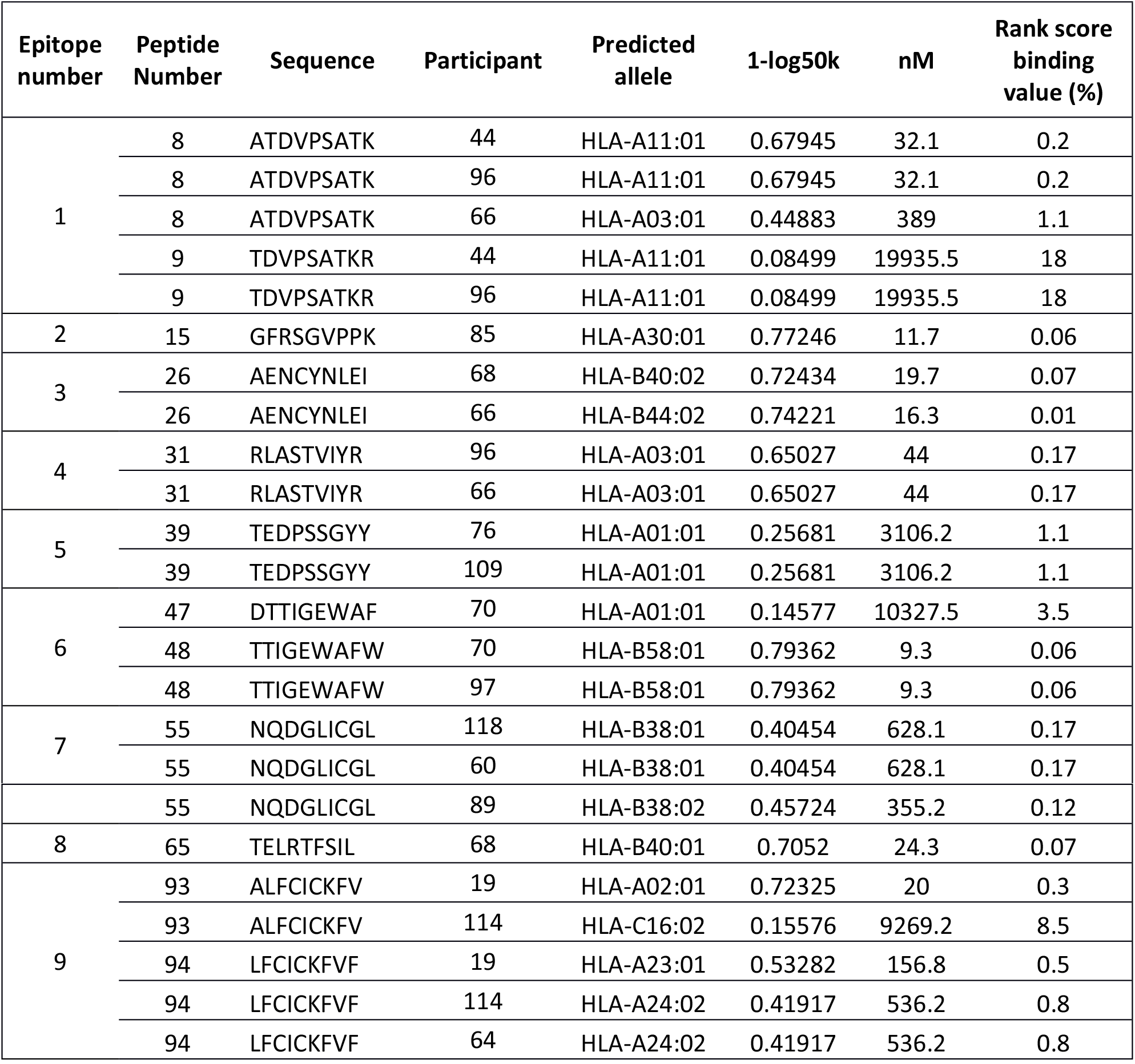
Predicted binding of epitopes to participant HLA alleles.

There were three epitopes where the prediction algorithm gave a poor (rank > 2%) binding prediction for the HLA of some of the responding volunteers. In each case, a shift of a single amino acid produced a strong binding value. In these cases, the assays were repeated and the result reconfirmed (data not shown). An example of this was the epitope TDVPSATKR. Here, 2 volunteers (44 and 96) expressed the HLA-A11:01 allele, yet this was predicted to bind very poorly (rank score 18%). The adjacent epitope ATDVPSATK was predicted to bind strongly with this allele and was indeed recognised by three volunteers expressing this allele.

To confirm recognition of epitopes through the predicted allele, MHC Class I pentamers were produced for peptide 1 (ATDFPSATK) bound to the HLA-A11:01 MHC molecule and a second for peptide 9 (LFCICKFVF) bound to HLA-A24:02. For the HLA-A11:01 pentamer containing the ATDVPSATK peptide, 0.83% and 0.15% of CD8^+^ T cells stained positive from donor 044 and 096 respectively, while PBMC from a mis-matched volunteer, 066, were negative (Figure 5A). Only one volunteer recognising LFCICKFVF had sufficient PBMC remaining for pentamer staining and 0.14% of their CD8^+^ T cells were pentamer positive (Figure 5B). We therefore demonstrated that the *in silico* prediction of MHC allele binding to epitopes we identified was correct for at least two nonamers.

**Figure 5.**
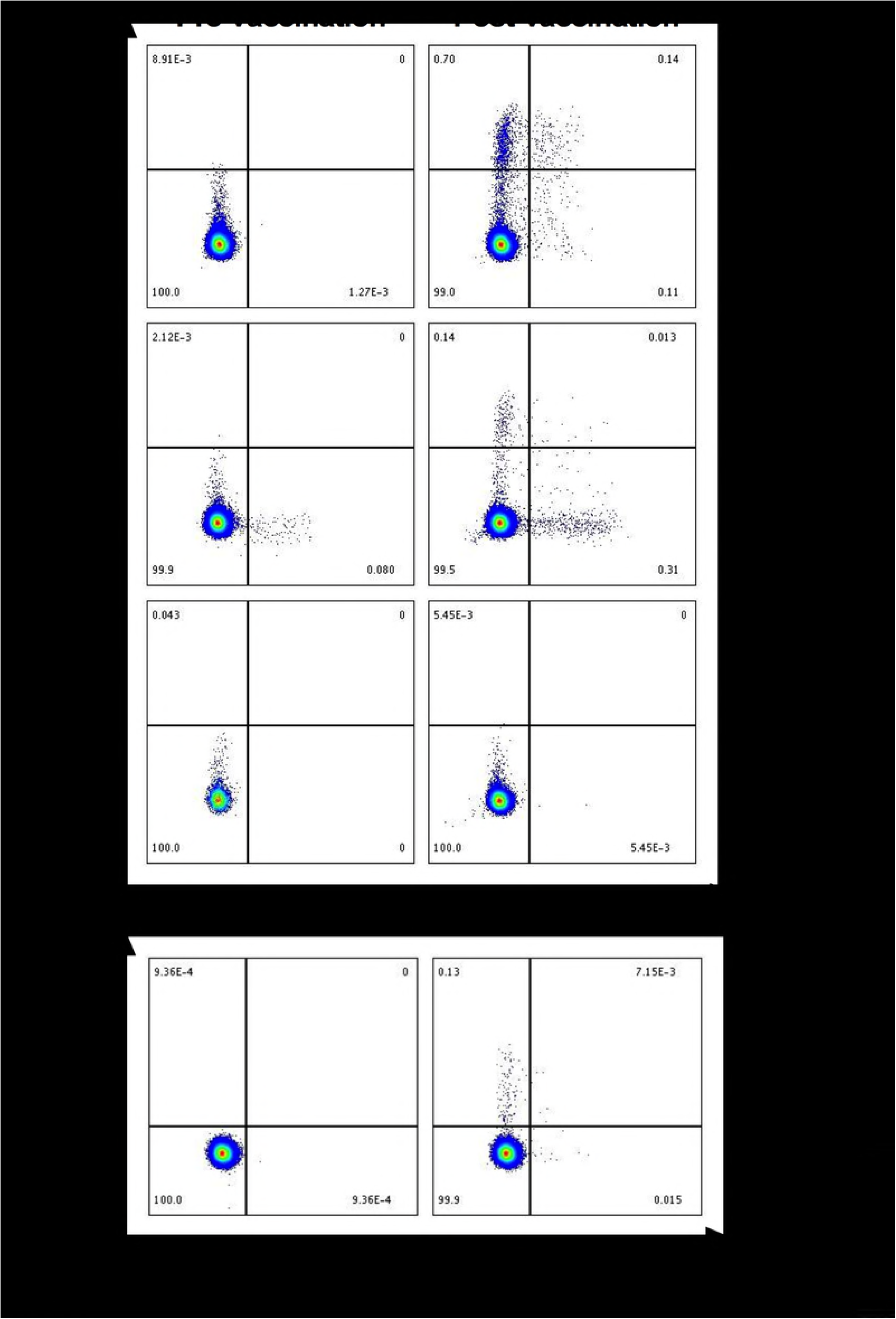
Intracellular cytokine staining with pentamer staining. (A) An HLA-A11:01 pentamer containing the ATDVPSATK peptide was used to stain PBMC from 3 volunteers, all shown to recognise this epitope (2 HLA-matches: 044 and 096, and 1 HLA-mismatch: 066) after stimulation with peptide. After subtraction of pre-vaccination background, the HLA-matched volunteers had pentamer positive populations of 0.83% and 0.15% of CD8^+^ T cells, respectively. No response was detected from the mismatched volunteer, 066. (B). PBMC from an HLA-matched volunteer were stained with an HLA-A24:02 pentamer containing the LFCICKFVF peptide. After subtraction of pre-vaccination background, 0.14% of CD8^+^ T cells were pentamer positive.

## Discussion

In this study, we identified 13 nonamers in the EBOV glycoprotein that elicit strong cellular responses in vaccinated individuals. When overlaid on to the protein, the nonamers form nine discreet epitopes, ranging in length from 9 to 14 amino acids (Supplementary Figure 1). Identifying the immunodominant regions of a protein builds our understanding of the immune response towards a pathogen and can help inform future vaccine design. Currently there are no licensed vaccines for Ebolavirus, however the unprecedented magnitude of the West African outbreak led to regulatory fast-tracking for several promising candidates. The most advanced of these vaccines is rVSV-ZEBOV which was assessed extensively in a Phase 3 clinical trial where high efficacy was demonstrated [44]. The platform for this vaccine is the zoonotic rhabdovirus vesicular stomatitis virus, with its native glycoprotein substituted for EBOV-GP. Significant reactogenicity to rVSV-ZEBOV has been observed in some populations, including fever and arthralgia in 25% and 22% of recipients in a Swiss cohort [45], perhaps due to the replication competency of the vector. In contrast, the ChAd3 and MVA vectors used in this study are replication-deficient and substantially less reactogenic, eliciting an immune response comprising both potent antibody and cellular components [46]. Induction of high frequency CD8^+^ T cell responses is critical to survival in NHP models of EBOV and has also now been demonstrated in human survivors of EVD, suggesting a potential advantage for these vectors [7, 31].

Conservation of epitopes is critical if they are to be of use in future vaccines as alterations to the T cell targets will likely result in diminished efficacy. Initial reports from the West African outbreak indicated that the circulating EBOV was undergoing an increased mutation rate, up to twice that seen in previous outbreaks [47]. Further work subsequently demonstrated that the viral mutation rate was in fact similar to those previously estimated with the majority of nucleotide alterations being either silent or occurring in noncoding regions [48, 49]. The epitopes identified here exhibit a high level of conservation across EBOV strains. The ChAd3 and MVA vectors encode GP from the 1976 Mayinga strain, and at the amino acid level, the epitopes are completely conserved in the 2014 Makona-Gueckedou strain as well as all intervening strains which were identified (Kikwit 1995, Gabon 1996, Gabon 2002). Most of these epitopes are highly conserved with the related SUDV GP, suggesting that T cells raised in individuals vaccinated with EBOV GP would show cross-reactivity against SUDV GP.

Interestingly, three of the epitopes characterised within this study had previously been predicted, in part or completely, by computational and murine studies. The fourth epitope was partially predicted by two different studies and shown to effectively stimulate splenocytes of immunised mice [28, 30, 50]. Epitopes seven and eight were also predicted, the former partially and the later entirely, by several studies [27–29]. We were also able to confirm the accuracy of the epitope prediction algorithm with pentamers for 2 epitopes, showing that *in silico* approaches remain a valid strategy to infer MHC restriction.

The epitopes identified in this paper are the first human EBOV T cell epitopes in the glycoprotein characterised in detail and build our knowledge about cellular responses to this pathogen. Identification and characterisation of the identified peptides may also aid in future vaccine design. Determining the strongly and weakly immunogenic regions of a target protein could allow vaccine constructs to be truncated or create mosaic inserts for vectors with a limited genomic capacity. Importantly, the finding that this vaccine regime elicits potent cell-mediated immunity in donors with a broad range of HLA types indicates the potential immunogenicity of this approach in genetically diverse populations. Furthermore, dominant epitopes induced by vaccination could be compared with those elicited by natural infection in survivors of Ebolavirus disease, thus allowing inference of potential protective ability of different vaccination strategies in lieu of data from an efficacy trial in an outbreak setting. This study is an important addition to our understanding of the immune response to Ebolavirus, highlighting the importance of both humoral and cellular responses during infection and vaccination.

## Materials and methods

### Study design

Healthy adult volunteers (n=60) were vaccinated with ChAd3-ZEBOV (1-5×10^10^ viral particles) as part of a recent clinical trial [ClinicalTrials.gov number, NCT02240875][33]. Of these, 30 subjects received vaccination with MVA-BN filo (1.5-3×10^8^ plaque-forming units) as a heterologous boost 1-10 weeks after the priming vaccination. The original clinical trial from which these samples were obtained was supported by the Wellcome Trust, the United Kingdom Medical Research Council, the United Kingdom Department for International Development, and the United Kingdom National Institute for Health Research Oxford Biomedical Research Centre (106325/Z/14/A). The clinical trial protocol was published with the original clinical study [33] and a CONSORT diagram and checklist are provided in the online Supplementary Appendix.

### Ethics statement

The study was reviewed and approved by the United Kingdom National Research Ethics Service, the Committee South Central–Oxford A, the Medicines and Healthcare Products Regulatory Agency, and the Oxford University Clinical Trials and Research Governance team, who monitored compliance with Good Clinical Practice guidelines. An independent data and safety monitoring board provided safety oversight. Written informed consent was obtained from all participants and all participants were adults. The study was conducted in compliance with the clinical trial protocol, International Conference on Harmonisation Good Clinical Practice Guideline E6 (R1) (ICH-GCP) and the applicable regulatory requirements.

### Blood processing

Blood samples were stored at room temperature prior to processing, which was completed within six hours of venepuncture. PBMC were separated by density centrifugation from heparinised whole blood and resuspended in R10 medium (RPMI containing 10% heat-inactivated, batch-tested, sterile-filtered foetal calf serum [FCS] previously screened for low reactivity [Labtech International], 1% L-glutamine, 1% penicillin/streptomycin). Cell counts were performed using a CASY Cell Counter (Roche Innovatis AG) according to an established SOP in the lab.

### *Ex vivo* IFNγ ELISpot assays

*Ex vivo* (16-18 hour stimulation) ELISpot assays were performed using Multiscreen IP ELISpot plates (Millipore), human IFNγ SA-ALP antibody kits (Mabtech) and BCIP NBT-plus chromogenic substrate (Moss Inc.). Initial ELISPOT assays were performed using freshly isolated PBMC as part of the immunomonitoring for the clinical trial (Figure 1A). All subsequent ELISpot assays were performed using PBMC cryopreserved from the clinical trial and stored in the vapour phase of liquid nitrogen. Vials of frozen PBMC were thawed in a 37°C water bath, washed in R10 and incubated at 1-5 million PBMC per ml of R10 for 2-4 hours with 250IU of benzonase per ml of R10. PBMC were tested in triplicate against antigens at 2.5μg/ml with 200,000 PBMC added to each well of the ELISpot plate. Plates were counted using an AID automated ELISpot counter (AID Diagnostika GmbH, algorithm C), using identical settings for all plates, and counts were adjusted only to remove artefacts. Responses were averaged across triplicate wells, responses in unstimulated (negative control) wells were subtracted. Staphylococcal Enterotoxin B (0.02μg/ml) and phytohaemmagglutinin-L (10μg/ml) were used as a positive control, whereby responses of >1000 SFC/10^6^ PBMC passed QC.

### Epitope Identification

From the initial study, a peptide library consisting of 187 peptides (12-17 amino acids in length, overlapping by 4 residues) spanning the entire EBOV glycoprotein was synthesised (Neobiolab, MA, USA). These were initially dissolved in DMSO to 100mg/ml, combined into 10 separate pools and diluted in PBS to 7.5μg/ml for the clinical trial ELISpots (Supplementary Table 1). The peak of the immune response after vaccination was seven days post-MVA and this time point was used to identify the most immunogenic peptide pools. A positive response to a peptide pool was arbitrarily defined as a response greater than four standard deviations of the median of the negative control response for all volunteers (equivalent to 78 SFC). Peptide pools that elicited positive responses in more than 40% of participants were selected for further epitope mapping. Three of the pools (SP, GP1.5 and GP1.7) were not studied further due to low responses. Samples from participants with the highest peak responses against the remaining seven peptide pools were selected (taking into account sample availability) and tested against the individual constituent peptides of the respective pools to identify immunodominant peptides (n=6-8 participants per pool, 26 in total). Responses were considered positive if they were greater than four standard deviations of the median of the negative control response for all volunteers (equivalent to 17 SFC for frozen assays). A selection of the individual peptides or region or peptides that elicited the highest responses were resynthesized as peptide libraries of nonamers overlapping by a single amino acid to identify the minimal epitope (ProImmune Ltd, Oxford, UK). These were tested using frozen PBMC as described above.

### Epitope Mapping

The EBOV GP domain organization scheme was produced by using DOG software, version 2.0 [51]. The crystal structure of EBOV GP (PDB ID 5JQ3 [52]) was visualized using the PyMOL Molecular Graphics System, Version 1.705 Schrödinger, LLC (https://pymol.org/). The footprints of the endosomal receptor Niemann-Pick C1 (NPC1) and human neutralizing antibody (nAb) KZ52 on the EBOV GP were obtained by analysing the corresponding complex crystal structures (PDB ID 5F1B [53] and PDB ID 3CSY [14]) using the ‘Protein interfaces, surfaces and assemblies’ service PISA at the European Bioinformatics Institute (http://www.ebi.ac.uk/pdbe/prot_int/pistart.html) [54].

### Intracellular Cytokine Staining (ICS)

Due to sample constraints, ICS was carried out on eight nonamers that elicited the highest responses by ELISpot (cut off >475 SFC/10^6^ PBMC). Frozen PBMC were thawed as described above and ICS was performed as described previously (34). Cells were stimulated with single peptides at 2.5μg/ml. A minimum of 1×10^6^ events were acquired on a LSR II cytometer (Becton Dickinson, Oxford, UK). Data were prepared and analysed using FlowJo v9.8.1 (Treestar Inc., Ashland, Oregon, USA). A hierarchical gating strategy was used. Responses to the peptides for each sample were determined after subtracting the responses from unstimulated controls.

### Pentamer staining and flow cytometry

R-PE-labelled custom Pro5^®^ pentamers (ProImmune, Oxford, UK) were synthesised for epitope 1 (ATDFPSATK) bound to the HLA-A11:01 allele and a second for epitope 9 (LFCICKFVF) bound to the HLA-A24:02 allele. PBMC from pre-vaccination and post-vaccination (Day 84) time points were stained and analysed by flow cytometry. After thawing, 10μL of the pentamer was added to each PBMC sample and incubated for 10 minutes at room temperature in the dark. After washing, samples were resuspended in media containing 1μg/mL of anti-CD28 and anti-CD49d and 1μL anti-CD107a. Peptides were added at a final concentration of 2μg/mL per peptide and samples incubated at 37°C. Brefeldin A and monensin were added after 2 hours, samples were then incubated at 37°C for a further 16 hours. Data were prepared and analysed using FlowJo v9.8.1 (Treestar Inc., Ashland, Oregon, USA). A hierarchical gating strategy was used. Responses to the peptides for each sample were determined after subtracting the responses from pre-vaccination samples. PBMC from a responder to the same epitope with mismatched alleles was also stained with the epitope 1 pentamer, however insufficient remaining PBMC were available for volunteers recognising epitope 9.

### Epitope HLA-restriction predication

MHC Class I alleles were analysed using NetMHCpan (version 3.0) to assign an HLA restriction element from each volunteer to each of the positive peptides. HLA restriction and peptide binding strength was assigned by for each peptide predicting binding to all HLA molecules of the positive volunteer and reporting the lowest percentile rank score and associated HLA molecule. Rank score binding values less than 2% were considered to predict strong binding of an epitope to a HLA allele. This is the value conventionally used to define peptide binders to HLA. At this binding value 95% of validated epitopes are identified at a specificity of > 99% [55].

## Acknowledgements

The ChAd3 vaccine was provided by the Vaccine Research Center of the National Institute of Allergy and Infectious Diseases (NIAID) and GlaxoSmithKline. MVA-BN Filo was produced under a contract (FBS-004-009) between the NIAID and Fisher BioServices and a contract (HHSN272200800044C) between the National Institutes of Health and Fisher Bioservices. The original clinical trial was supported by the Welcome Trust (106325/Z/14/Z) and additional support was provided by by the National Institute for Health Research (NIHR) Oxford Biomedical Research Centre (BRC). The views expressed are those of the author(s) and not necessarily those of the NHS, the NIHR or the Department of Health. We are grateful to all of the clinicians, nurses, scientists, project managers and investigators who were involved in the clinical trial that generated the samples used in this study. HLA-typing was performed by the Transplant Immunology & Immunogenetics laboratory of the Oxford University Hospitals NHS Trust. TAB is supported by the MRC (MR/L009528/1) and the Wellcome Centre for Human Genetics is supported by grant 203141/Z/16/Z. We thank Professor Miles Carroll, Dr Yper Hall and Dr Tom Tipton of Public Health England for helpful discussions and Dr Sarah Sebastian for reviewing the manuscript.

## Supporting Information Legends

Supplementary Table 1. Amino acid sequences of overlapping peptides spanning the length of the ZEBOV glycoprotein, a.a.-amino acids

Supplementary Table 2. Amino acid sequences of nonamer peptides used for mapping immunodominant peptides within peptide pools, a.a.-amino acids. Peptide relates to peptide number in Supplementary Table 1.

Supplementary Table 3. HLA types for HLA-A, B and C alleles for volunteers whose samples were used in epitope mapping experiments. A dash indicates homozygous alleles. ELISpot response is that measured in the initial assay using 15mer pools.

**Supplementary Figure 1.** Mapping the identified T cell epitopes onto Ebola virus glycoprotein (EBOV GP) (A) Location of the T cell epitopes is mapped onto the crystal structure of EBOV GP (PDB [Protein Data Bank] ID 5JQ3 (36)) using the same colour scheme as in Figure 3. The T cell epitopes 206-214 and 282-291 are not fully visualized due to crystallographically disordered residues 206-210, 285 and 286. T cell epitope 667-676 is not included in the crystallized construct. (B) Location of the T cell epitopes visualized after removal of the glycan cap from EBOV GP. Footprints of the endosomal receptor Niemann-Pick C1 (NPC1) and human neutralizing antibody (nAb) KZ52 are shown in green. The overlaps between the T cell epitopes and the NPC1 or KZ52 binding sites are emphasized in dark green. For clarity, each binding site is shown on one of the three monomers only.

**Supplementary Figure 2**. Flow cytometry gating strategy. Singlets were identified using forward scatter plots. Dead cells were excluded by violet fluorescent amine-reactive dye staining. Monocytes and B cells were excluded by CD14 or CD19 expression and T cells identified by CD3 expression. T cells were then subdivided by gating on CD4+ and CD8^+^ populations. Cytokine expression was quantified by plotting pairs of cytokines against each other and gating positive populations. This is a representative sample from a sample stimulated overnight (18 hours) with a single pool of overlapping GP peptides.

**Supplementary Figure 3. Flow cytometry gating strategy for pentamer staining**. Singlets were identified using forward scatter plots. Dead cells were excluded by violet fluorescent amine-reactive dye staining. Monocytes and B cells were excluded by CD14 or CD19 expression and T cells identified by CD3 expression. CD8 positive and CD4 negative cells were then selected, then IFNγ expression and pentamer binding were determined.

## Bibliography

1. WHO. Ebola Situation Report-5th June 2017. External Situation Report 21 [Internet]. 2017 9th June 2017; (5th June 2017):[1–6 pp.]. Available from: http://apps.who.int/iris/bitstream/10665/255630/1/EbolaDRC-06062017.pdf?ua=1.

2. Baize S, Pannetier D, Oestereich L, Rieger T, Koivogui L, Magassouba N, et al. Emergence of Zaire Ebola virus disease in Guinea. N Engl J Med. 2014;371(15):1418–25. Epub 2014/04/18. doi: 10.1056/NEJMoa1404505. PubMed PMID: 24738640.

3. Kuhn JH, Becker S, Ebihara H, Geisbert TW, Johnson KM, Kawaoka Y, et al. Proposal for a revised taxonomy of the family Filoviridae: classification, names of taxa and viruses, and virus abbreviations. Archives of virology. 2010;155(12):2083–103. doi: 10.1007/s00705-010-0814-x. PubMed PMID: PMC3074192.

4. Feldmann H, Geisbert TW. Ebola haemorrhagic fever. Lancet. 2011;377(9768):849–62. Epub 2010/11/19. doi: 10.1016/S0140-6736(10)60667-8. PubMed PMID: 21084112; PubMed Central PMCID: PMCPMC3406178.

5. Jahrling PB, Geisbert TW, Dalgard DW, Johnson ED, Ksiazek TG, Hall WC, et al. Preliminary report: isolation of Ebola virus from monkeys imported to USA. Lancet. 1990;335(8688):502–5. Epub 1990/03/03. PubMed PMID: 1968529.

6. Barrette RW, Metwally SA, Rowland JM, Xu L, Zaki SR, Nichol ST, et al. Discovery of swine as a host for the Reston ebolavirus. Science. 2009;325(5937):204–6. Epub 2009/07/11. doi: 10.1126/science.1172705. PubMed PMID: 19590002.

7. Sullivan NJ, Martin JE, Graham BS, Nabel GJ. Correlates of protective immunity for Ebola vaccines: implications for regulatory approval by the animal rule. Nature reviews Microbiology. 2009;7(5):393–400. Epub 2009/04/17. doi: 10.1038/nrmicro2129. PubMed PMID: 19369954.

8. Dahlke C, Lunemann S, Kasonta R, Kreuels B, Schmiedel S, Ly ML, et al. Comprehensive Characterization of Cellular Immune Responses Following Ebola Virus Infection. J Infect Dis. 2017;215(2):287–92. Epub 2016/11/02. doi: 10.1093/infdis/jiw508. PubMed PMID: 27799354.

9. McElroy AK, Akondy RS, Davis CW, Ellebedy AH, Mehta AK, Kraft CS, et al. Human Ebola virus infection results in substantial immune activation. Proc Natl Acad Sci U S A. 2015;112(15):4719–24. Epub 2015/03/17. doi: 10.1073/pnas.1502619112. PubMed PMID: 25775592; PubMed Central PMCID: PMC4403189.

10. Lee JE, Saphire EO. Neutralizing ebolavirus: structural insights into the envelope glycoprotein and antibodies targeted against it. Curr Opin Struct Biol. 2009;19(4):408–17. Epub 2009/06/30. doi: 10.1016/j.sbi.2009.05.004. PubMed PMID: 19559599; PubMed Central PMCID: PMCPMC2759674.

11. Ruibal P, Oestereich L, Ludtke A, Becker-Ziaja B, Wozniak DM, Kerber R, et al. Unique human immune signature of Ebola virus disease in Guinea. Nature. 2016;533(7601):100–4. Epub 2016/05/07. doi: 10.1038/nature17949. PubMed PMID: 27147028; PubMed Central PMCID: PMCPMC4876960.

12. Becquart P, Mahlakoiv T, Nkoghe D, Leroy EM. Identification of continuous human B-cell epitopes in the VP35, VP40, nucleoprotein and glycoprotein of Ebola virus. PLoS One. 2014;9(6):e96360. Epub 2014/06/11. doi: 10.1371/journal.pone.0096360. PubMed PMID: 24914933; PubMed Central PMCID: PMCPMC4051576.

13. Lee JE, Saphire EO. Ebolavirus glycoprotein structure and mechanism of entry. Future Virol. 2009;4(6):621–35. Epub 2009/01/01. doi: 10.2217/fvl.09.56. PubMed PMID: 20198110; PubMed Central PMCID: PMCPMC2829775.

14. Lee JE, Fusco ML, Hessell AJ, Oswald WB, Burton DR, Saphire EO. Structure of the Ebola virus glycoprotein bound to an antibody from a human survivor. Nature. 2008;454(7201):177–82. Epub 2008/07/11. doi: 10.1038/nature07082. PubMed PMID: 18615077; PubMed Central PMCID: PMCPMC2700032.

15. Corti D, Misasi J, Mulangu S, Stanley DA, Kanekiyo M, Wollen S, et al. Protective monotherapy against lethal Ebola virus infection by a potently neutralizing antibody. Science. 2016;351(6279):1339–42. Epub 2016/02/27. doi: 10.1126/science.aad5224. PubMed PMID: 26917593.

16. Parren PW, Geisbert TW, Maruyama T, Jahrling PB, Burton DR. Pre- and postexposure prophylaxis of Ebola virus infection in an animal model by passive transfer of a neutralizing human antibody. J Virol. 2002;76(12):6408–12. Epub 2002/05/22. PubMed PMID: 12021376; PubMed Central PMCID: PMCPMC136210.

17. Qiu X, Wong G, Audet J, Bello A, Fernando L, Alimonti JB, et al. Reversion of advanced Ebola virus disease in nonhuman primates with ZMapp. Nature. 2014;514(7520):47–53. Epub 2014/08/30. doi: 10.1038/nature13777. PubMed PMID: 25171469; PubMed Central PMCID: PMCPMC4214273.

18. Pettitt J, Zeitlin L, Kim DH, Working C, Johnson JC, Bohorov O, et al. Therapeutic intervention of Ebola virus infection in rhesus macaques with the MB-003 monoclonal antibody cocktail. Sci Transl Med. 2013;5(199):199ra13. Epub 2013/08/24. doi: 10.1126/scitranslmed.3006608. PubMed PMID: 23966302.

19. Henao-Restrepo AM, Longini IM, Egger M, Dean NE, Edmunds WJ, Camacho A, et al. Efficacy and effectiveness of an rVSV-vectored vaccine expressing Ebola surface glycoprotein: interim results from the Guinea ring vaccination cluster-randomised trial. Lancet. 2015;386(9996):857–66. Epub 2015/08/08. doi: 10.1016/S0140-6736(15)61117-5. PubMed PMID: 26248676.

20. Agnandji ST, Huttner A, Zinser ME, Njuguna P, Dahlke C, Fernandes JF, et al. Phase 1 Trials of rVSV Ebola Vaccine in Africa and Europe. N Engl J Med. 2016;374(17):1647–60. Epub 2015/04/02. doi: 10.1056/NEJMoa1502924. PubMed PMID: 25830326; PubMed Central PMCID: PMCPMC5490784.

21. Rechtien A, Richert L, Lorenzo H, Martrus G, Hejblum B, Dahlke C, et al. Systems Vaccinology Identifies an Early Innate Immune Signature as a Correlate of Antibody Responses to the Ebola Vaccine rVSV-ZEBOV. Cell Rep. 2017;20(9):2251–61. Epub 2017/08/31. doi: 10.1016/j.celrep.2017.08.023. PubMed PMID: 28854372; PubMed Central PMCID: PMCPMC5583508.

22. Dahlke C, Kasonta R, Lunemann S, Krahling V, Zinser ME, Biedenkopf N, et al. Dose-dependent T-cell Dynamics and Cytokine Cascade Following rVSV-ZEBOV Immunization. EBioMedicine. 2017;19:107–18. Epub 2017/04/25. doi: 10.1016/j.ebiom.2017.03.045. PubMed PMID: 28434944; PubMed Central PMCID: PMCPMC5440606.

23. Murin CD, Fusco ML, Bornholdt ZA, Qiu X, Olinger GG, Zeitlin L, et al. Structures of protective antibodies reveal sites of vulnerability on Ebola virus. Proc Natl Acad Sci U S A. 2014;111(48):17182–7. Epub 2014/11/19. doi: 10.1073/pnas.1414164111. PubMed PMID: 25404321; PubMed Central PMCID: PMCPMC4260551.

24. Misasi J, Gilman MS, Kanekiyo M, Gui M, Cagigi A, Mulangu S, et al. Structural and molecular basis for Ebola virus neutralization by protective human antibodies. Science. 2016;351(6279):1343–6. Epub 2016/02/27. doi: 10.1126/science.aad6117. PubMed PMID: 26917592; PubMed Central PMCID: PMCPMC5241105.

25. Sullivan NJ, Hensley L, Asiedu C, Geisbert TW, Stanley D, Johnson J, et al. CD8^+^ cellular immunity mediates rAd5 vaccine protection against Ebola virus infection of nonhuman primates. Nat Med. 2011;17(9):1128–31. Epub 2011/08/23. doi: 10.1038/nm.2447. PubMed PMID: 21857654.

26. Theaker SM, Rius C, Greenshields-Watson A, Lloyd A, Trimby A, Fuller A, et al. T-cell libraries allow simple parallel generation of multiple peptide-specific human T-cell clones. J Immunol Methods. 2016;430:43–50. Epub 2016/01/31. doi: 10.1016/j.jim.2016.01.014. PubMed PMID: 26826277; PubMed Central PMCID: PMCPMC4783706.

27. Simmons G, Lee A, Rennekamp AJ, Fan X, Bates P, Shen H. Identification of murine T-cell epitopes in Ebola virus nucleoprotein. Virology. 2004;318(1):224–30. Epub 2004/02/20. doi: 10.1016/j.virol.2003.09.016. PubMed PMID: 14972550.

28. Dikhit MR, Kumar S, Vijay mahantesh, Sahoo BR, Mansuri R, Amit A, et al. Computational elucidation of potential antigenic CTL epitopes in Ebola virus. Infect Genet Evol. 2015;36:369–75. Epub 2015/10/16. doi: 10.1016/j.meegid.2015.10.012. PubMed PMID: 26462623.

29. Sundar K, Boesen A, Coico R. Computational prediction and identification of HLA-A2.1-specific Ebola virus CTL epitopes. Virology. 2007;360(2):257–63. Epub 2006/11/25. doi: 10.1016/j.virol.2006.09.042. PubMed PMID: 17123567.

30. Yasmin T, Nabi AHMN. B and T Cell Epitope-Based Peptides Predicted from Evolutionarily Conserved and Whole Protein Sequences of Ebola Virus as Vaccine Targets. Scand J Immunol. 2016;83(5):321–37. doi: 10.1111/sji.12425.

31. Sakabe S, Sullivan BM, Hartnett JN, Robles-Sikisaka R, Gangavarapu K, Cubitt B, et al. Analysis of CD8(+) T cell response during the 2013-2016 Ebola epidemic in West Africa. Proc Natl Acad Sci U S A. 2018;115(32):E7578–E86. Epub 2018/07/25. doi: 10.1073/pnas.1806200115. PubMed PMID: 30038008; PubMed Central PMCID: PMCPMC6094108.

32. Milligan ID, Gibani MM, Sewell R, Clutterbuck EA, Campbell D, Plested E, et al. Safety and Immunogenicity of Novel Adenovirus Type 26- and Modified Vaccinia Ankara-Vectored Ebola Vaccines: A Randomized Clinical Trial. JAMA. 2016;315(15):1610–23. doi: 10.1001/jama.2016.4218. PubMed PMID: 27092831.

33. Ewer K, Rampling T, Venkatraman N, Bowyer G, Wright D, Lambe T, et al. A Monovalent Chimpanzee Adenovirus Ebola Vaccine Boosted with MVA. N Engl J Med. 2016;374(17):1635–46. Epub 2015/01/30. doi: 10.1056/NEJMoa1411627. PubMed PMID: 25629663; PubMed Central PMCID: PMCPMC5798586.

34. Venkatraman N, Ndiaye BP, Bowyer G, Wade D, Sridhar S, Wright D, et al. Safety and immunogenicity of a heterologous prime-boost Ebola virus vaccine regimen - ChAd3-EBO-Z followed by MVA-EBO-Z in healthy adults in the UK and Senegal. J Infect Dis. 2018. Epub 2018/11/09. doi: 10.1093/infdis/jiy639. PubMed PMID: 30407513.

35. Sanchez-Trincado JL, Gomez-Perosanz M, Reche PA. Fundamentals and Methods for T- and B-Cell Epitope Prediction. J Immunol Res. 2017;2017:2680160. Epub 2018/02/16. doi: 10.1155/2017/2680160. PubMed PMID: 29445754; PubMed Central PMCID: PMCPMC5763123.

36. Soria-Guerra RE, Nieto-Gomez R, Govea-Alonso DO, Rosales-Mendoza S. An overview of bioinformatics tools for epitope prediction: implications on vaccine development. J Biomed Inform. 2015;53:405–14. Epub 2014/12/03. doi: 10.1016/j.jbi.2014.11.003. PubMed PMID: 25464113.

37. Hoof I, Peters B, Sidney J, Pedersen LE, Sette A, Lund O, et al. NetMHCpan, a method for MHC class I binding prediction beyond humans. Immunogenetics. 2009;61(1):1–13. Epub 2008/11/13. doi: 10.1007/s00251-008-0341-z. PubMed PMID: 19002680; PubMed Central PMCID: PMCPMC3319061.

38. Azizi A, Anderson DE, Torres JV, Ogrel A, Ghorbani M, Soare C, et al. Induction of broad cross-subtype-specific HIV-1 immune responses by a novel multivalent HIV-1 peptide vaccine in cynomolgus macaques. J Immunol. 2008;180(4):2174–86. Epub 2008/02/06. PubMed PMID: 18250424.

39. Fischer W, Perkins S, Theiler J, Bhattacharya T, Yusim K, Funkhouser R, et al. Polyvalent vaccines for optimal coverage of potential T-cell epitopes in global HIV-1 variants. Nat Med. 2007;13(1):100–6. Epub 2006/12/26. doi: 10.1038/nm1461. PubMed PMID: 17187074.

40. Ondondo B, Murakoshi H, Clutton G, Abdul-Jawad S, Wee EG, Gatanaga H, et al. Novel Conserved-region T-cell Mosaic Vaccine With High Global HIV-1 Coverage Is Recognized by Protective Responses in Untreated Infection. Mol Ther. 2016;24(4):832–42. Epub 2016/01/09. doi: 10.1038/mt.2016.3. PubMed PMID: 26743582; PubMed Central PMCID: PMCPMC4886941.

41. McMichael AJ, Gotch FM, Noble GR, Beare PA. Cytotoxic T-cell immunity to influenza. N Engl J Med. 1983;309(1):13–7. Epub 1983/07/07. doi: 10.1056/NEJM198307073090103. PubMed PMID: 6602294.

42. Wu C, Zanker D, Valkenburg S, Tan B, Kedzierska K, Zou QM, et al. Systematic identification of immunodominant CD8^+^ T-cell responses to influenza A virus in HLA-A2 individuals. Proc Natl Acad Sci U S A. 2011;108(22):9178–83. Epub 2011/05/13. doi: 10.1073/pnas.1105624108. PubMed PMID: 21562214; PubMed Central PMCID: PMCPMC3107317.

43. Gras S, Kedzierski L, Valkenburg SA, Laurie K, Liu YC, Denholm JT, et al. Cross-reactive CD8^+^ T-cell immunity between the pandemic H1N1-2009 and H1N1-1918 influenza A viruses. Proc Natl Acad Sci U S A. 2010;107(28):12599–604. Epub 2010/07/10. doi: 10.1073/pnas.1007270107. PubMed PMID: 20616031; PubMed Central PMCID: PMCPMC2906563.

44. Henao-Restrepo AM, Camacho A, Longini IM, Watson CH, Edmunds WJ, Egger M, et al. Efficacy and effectiveness of an rVSV-vectored vaccine in preventing Ebola virus disease: final results from the Guinea ring vaccination, open-label, cluster-randomised trial (Ebola Ca Suffit!). Lancet. 2017;389(10068):505–18. Epub 2016/12/27. doi: 10.1016/S0140-6736(16)32621-6. PubMed PMID: 28017403; PubMed Central PMCID: PMCPMC5364328.

45. Huttner A, Dayer JA, Yerly S, Combescure C, Auderset F, Desmeules J, et al. The effect of dose on the safety and immunogenicity of the VSV Ebola candidate vaccine: a randomised double-blind, placebo-controlled phase 1/2 trial. Lancet Infect Dis. 2015;15(10):1156–66. doi: 10.1016/S1473-3099(15)00154-1. PubMed PMID: 26248510.

46. Ewer K, Rampling T, Venkatraman N, Bowyer G, Wright D, Lambe T, et al. A Monovalent Chimpanzee Adenovirus Ebola Vaccine Boosted with MVA. New England Journal of Medicine. 2016;374(17):1635–46. doi: doi:10.1056/NEJMoa1411627. PubMed PMID: 25629663.

47. Gire SK, Goba A, Andersen KG, Sealfon RS, Park DJ, Kanneh L, et al. Genomic surveillance elucidates Ebola virus origin and transmission during the 2014 outbreak. Science. 2014;345(6202):1369–72. Epub 2014/09/13. doi: 10.1126/science.1259657. PubMed PMID: 25214632; PubMed Central PMCID: PMCPMC4431643.

48. Olabode AS, Jiang X, Robertson DL, Lovell SC. Ebolavirus is evolving but not changing: No evidence for functional change in EBOV from 1976 to the 2014 outbreak. Virology. 2015;482:202–7. Epub 2015/04/17. doi: 10.1016/j.virol.2015.03.029. PubMed PMID: 25880111; PubMed Central PMCID: PMCPMC4503884.

49. Hoenen T, Safronetz D, Groseth A, Wollenberg KR, Koita OA, Diarra B, et al. Mutation rate and genotype variation of Ebola virus from Mali case sequences. Science. 2015;348(6230):117–9. doi: 10.1126/science.aaa5646.

50. Wu S, Yu T, Song X, Yi S, Hou L, Chen W. Prediction and identification of mouse cytotoxic T lymphocyte epitopes in Ebola virus glycoproteins. Virol J. 2012;9:111. Epub 2012/06/15. doi: 10.1186/1743-422X-9-111. PubMed PMID: 22695180; PubMed Central PMCID: PMCPMC3411508.

51. Ren J, Wen L, Gao X, Jin C, Xue Y, Yao X. DOG 1.0: illustrator of protein domain structures. Cell Res. 2009;19(2):271–3. Epub 2009/01/21. doi: 10.1038/cr.2009.6. PubMed PMID: 19153597.

52. Zhao Y, Ren J, Harlos K, Jones DM, Zeltina A, Bowden TA, et al. Toremifene interacts with and destabilizes the Ebola virus glycoprotein. Nature. 2016;535(7610):169–72. Epub 2016/07/01. doi: 10.1038/nature18615. PubMed PMID: 27362232; PubMed Central PMCID: PMCPMC4947387.

53. Wang H, Shi Y, Song J, Qi J, Lu G, Yan J, et al. Ebola Viral Glycoprotein Bound to Its Endosomal Receptor Niemann-Pick C1. Cell. 2016;164(1-2):258–68. doi: 10.1016/j.cell.2015.12.044. PubMed PMID: 26771495.

54. Krissinel E, Henrick K. Inference of macromolecular assemblies from crystalline state. J Mol Biol. 2007;372(3):774–97. Epub 2007/08/08. doi: 10.1016/j.jmb.2007.05.022. PubMed PMID: 17681537.

55. Lundegaard C, Lund O, Nielsen M. Accurate approximation method for prediction of class I MHC affinities for peptides of length 8, 10 and 11 using prediction tools trained on 9mers. Bioinformatics. 2008;24(11):1397–8. doi: 10.1093/bioinformatics/btn128.

